# A theta rhythm in macaque visual cortex and its attentional modulation

**DOI:** 10.1101/117804

**Authors:** Georgios Spyropoulos, Conrado A. Bosman, Pascal Fries

**Affiliations:** Ernst Strüngmann Institute (ESI) for Neuroscience in Cooperation with Max Planck Society, Deutschordenstraße 46, 60528 Frankfurt, Germany; Donders Institute for Brain, Cognition and Behaviour, Radboud University Nijmegen, Kapittelweg 29, 6525 EN Nijmegen, Netherlands; Swammerdam Institute for Life Sciences, Center for Neuroscience, Faculty of Science, University of Amsterdam, Sciencepark 904, 1098 XH Amsterdam, Netherlands

## Abstract

Theta rhythms govern rodent sniffing and whisking, and human language processing. Human psychophysics suggests a role for theta also in visual attention. Yet, little is known about theta in visual areas and its attentional modulation. We used electrocorticography (ECoG) to record local field potentials (LFPs) simultaneously from areas V1, V2, V4 and TEO of two macaque monkeys performing a selective visual attention task. We found a ≈4 Hz theta rhythm within both the V1-V2 and the V4-TEO region, and theta synchronization between them, with a predominantly feedforward directed influence. ECoG coverage of large parts of these regions revealed a surprising spatial correspondence between theta and visually induced gamma. Furthermore, gamma power was modulated with theta phase. Selective attention to the respective visual stimulus strongly reduced these theta-rhythmic processes, leading to an unusually strong attention effect for V1. Microsaccades (MSs) were partly locked to theta. Yet, neuronal theta rhythms tended to be even more pronounced for epochs devoid of MSs. Thus, we find an MS-independent theta rhythm specific to visually driven parts of V1-V2, which rhythmically modulates local gamma and entrains V4-TEO, and which is strongly reduced by attention. We propose that the less theta-rhythmic and thereby more continuous processing of the attended stimulus serves the exploitation of this behaviorally most relevant information. The theta-rhythmic and thereby intermittent processing of the unattended stimulus likely reflects the ecologically important exploration of less relevant sources of information.

## Introduction

In the human auditory system, a neuronal theta rhythm can lock to theta rhythmicity in speech signals and thereby support speech recognition (1). In rodent hippocampus, theta is particularly strong during navigational exploration of the environment (2, 3), and theta-related sequences of place cell firing seem to reflect the mental exploration of navigational options (4). In the rodent olfactory and somatosensory systems, sniffing and whisking often occur at a theta rhythm, exemplifying the role of theta in rhythmic sampling (5). Theta might serve a similar sampling function in the visual system. When human subjects freely view natural scenes, they perform saccades at a theta rhythm (6, 7). Saccades are overt expressions of shifts in visual spatial attention. Thus, visual attention might not be a continuous process, but rather explore visual input through theta rhythmic sampling. This seems to hold even in the absence of overt eye movements, for covert attentional sampling, as recently demonstrated through magnetoencephalography (MEG) (8). MEG shows visually induced gamma-band activity (GBA) (9) that is enhanced when the inducing stimulus is attended (10). Correspondingly, fluctuations in attention between two stimuli are reflected in the difference between their GBAs. This GBA difference reveals a theta rhythm that predicts change detection performance and thereby indexes covert attention (8).

This theta-rhythmic attentional sampling can even be observed directly in theta-rhythmic fluctuations of detection performance. When a human subject monitors two visual stimuli simultaneously, and a task-irrelevant flash is placed next to one of the stimuli, it draws attention and resets the rhythm. Subsequent randomly-timed changes in either one of the stimuli can reveal attentional allocation with high temporal resolution. The resulting time courses of detection performance show fluctuations of covert attention with a clear theta rhythmicity (11, 12). A related observation has also been described in inferotemporal (IT) cortex. In IT cortex, neuronal firing rates represent primarily the attended stimulus (13). When a single stimulus is presented, IT neurons respond with firing rates characteristic for the given stimulus. When a second stimulus is added to the screen, firing rates start oscillating at ≈4 Hz in a way that suggests that attention is drawn to the newly presented stimulus and subsequently alternates between the two stimuli (14). When a single stimulus is shown in isolation and receives full attention, IT neuronal firing rates show much weaker theta rhythmicity. These results suggest that attended stimuli might be processed in a more sustained manner, whereas stimuli that need to share attention with other stimuli are sampled theta rhythmically.

Attentional effects on firing rates in high-level ventral visual areas, like IT cortex, are subserved by the selective routing of attended stimuli through local and inter-areal gamma-band synchronization in lower ventral visual stream areas (7). Therefore, we hypothesized that lower ventral visual stream areas show a theta rhythm and theta-rhythmic modulations of gamma-band activity. We recorded local field potentials (LFPs) using high-density micro-electrocorticographic grids (ECoGs) simultaneously covering awake macaque areas V1, V2, V4 and TEO. We found ≈4 Hz theta-band activity within and synchronization between the post-lunate V1-V2 region and the pre-lunate V4-TEO region. The ECoG covered substantial parts of these areas, including both, visually driven and non-driven portions. This revealed that the portions showing a clear ≈4 Hz theta rhythm were very similar to the portions showing visually induced gamma-band activity. Furthermore, gamma amplitude was modulated by theta phase, that is, there was theta-gamma phase-amplitude coupling. Granger causality analysis revealed a stronger theta-rhythmic influence in the feedforward direction, from V1-V2 to V4-TEO, than in the feedback direction. Microsaccades were partly locked to the theta rhythm, yet, they did not explain the theta-rhythmic neuronal activity, because the rhythm tended to be even more pronounced in epochs devoid of microsaccades. Consistent with the notion that attended stimuli are processed more continuously, we found that selective attention to a visual stimulus reduced theta rhythmicity. The attention effects on theta rhythmicity and theta-gamma coupling were surprisingly strong for primary visual area V1.

## Results

### Macaque visual cortex shows a theta rhythm

We first calculated power spectra averaged over all electrodes on V1, V2, V4 and TEO from periods, during which the monkey fixated and covertly monitored either one of the two simultaneously presented drifting grating stimuli. Those average power spectra exhibited clear peaks in the gamma and the beta range, with peak frequencies specific to each monkey; however, they did not exhibit clear peaks in the theta-frequency range (Fig. 1A,B). We have previously found that power spectra can fail to reveal rhythms that are nevertheless unequivocally detectable with metrics of phase locking (15, 16). We therefore investigated phase locking in the low-frequency range. Phase locking metrics can be artificially inflated by a common recording reference, and therefore, we removed the recording reference by calculating local bipolar derivations, which we refer to as “sites”. We quantified phase locking by means of the pairwise phase consistency metric (PPC) (17). We calculated PPC spectra averaged over all possible pairs of sites within and between V1, V2, V4 and TEO. The average PPC spectra confirmed the gamma and beta peaks and in addition revealed clear theta peaks at 4-5 Hz (Fig. 1C,D). Thus, awake macaque visual cortex shows a distinct theta rhythm.

**Fig. 1.**
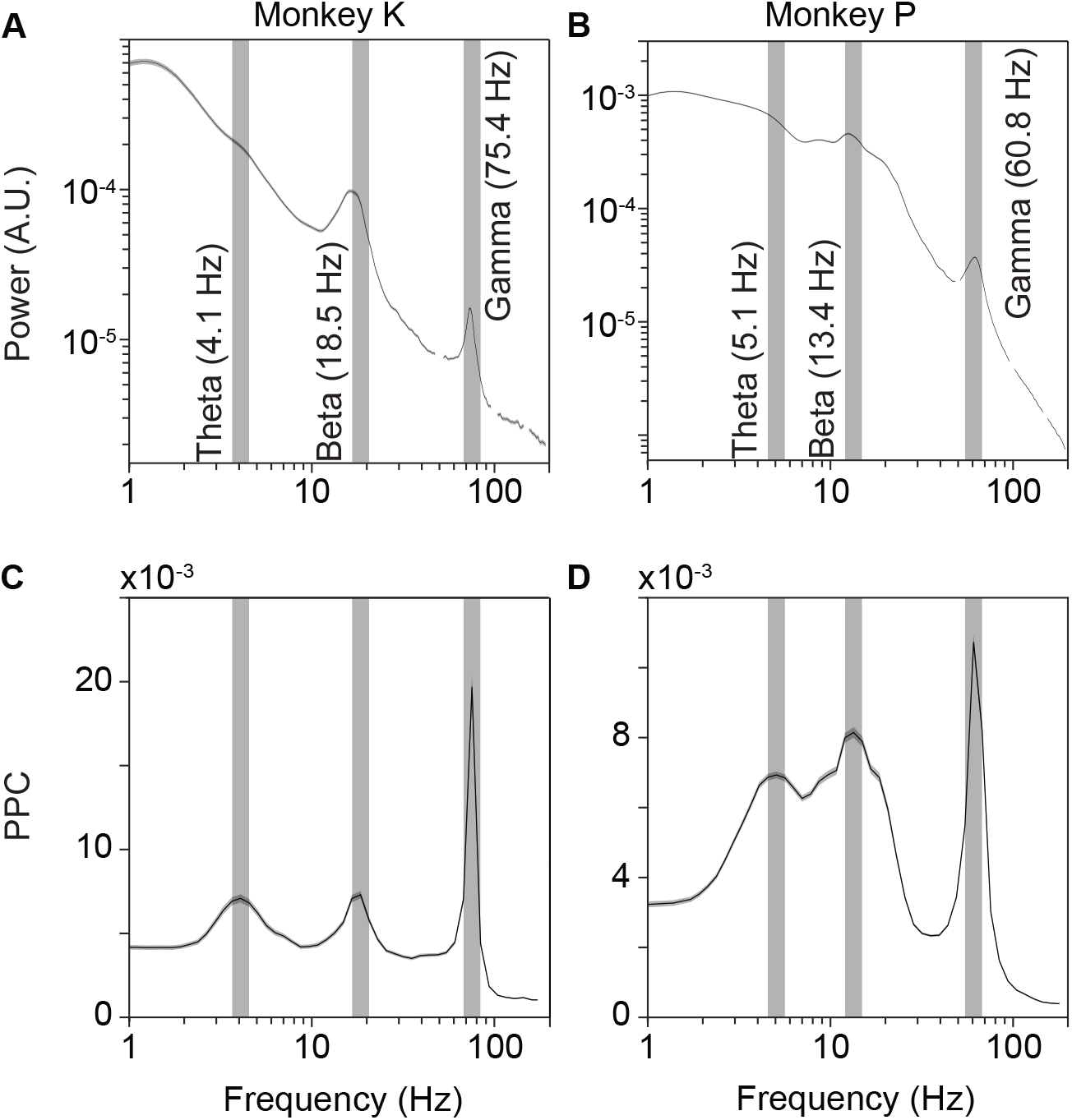
Average power and phase locking spectra for the two macaques. (*A*) Average power spectrum of the ECoG LFP in V1, V2, V4 and TEO, for monkey K. Data around the 50 Hz line-noise frequency and harmonics is not shown. (*B*) Same as *A*, but for monkey P. (*C*) Average phase locking (PPC) spectrum across all possible site pairs within and between V1, V2, V4 and TEO, for monkey K. (*D*), Same as C, but for monkey P. The error regions (hardly visible for *A* and *B*) show the 95% confidence interval based on a bootstrap procedure across data epochs. Vertical gray bars are shown at the peaks of the two macaques’ respective PPC spectra and are copied into *A* and *B*.

Note that the theta peaks in PPC spectra were not due to bipolar derivation. As we show below, some electrodes (without bipolar derivation) exhibited clear theta peaks in their power spectra, and those were actually reduced by calculating bipolar derivations. Thus phase-locking analysis revealed theta not because but despite bipolar derivation. Note also that the theta peaks in the PPC spectra were distinct from the shallow peaks in the power spectra between 1 and 2 Hz. These shallow power peaks were likely due to a reduction of power at the low end of the spectrum by the high-pass filter of the data acquisition system.

### Theta is spatially coextensive with visually induced gamma

We next investigated whether theta was related to visually induced activity and therefore first turned to the early visual areas covered by the ECoG grid. Area V1 extends from the posterior end of the brain towards the lunate sulcus, and V2 typically begins just posterior to the lunate. Thus, a few ECoG electrodes close to the lunate likely were on V2, and the rest of post-lunate ECoG electrodes were on V1. These ECoG electrodes provide gamma-power enhancements that are selective for particular stimulus positions, that is, gamma-power enhancements with circumscribed receptive fields (RFs) (18, 19). RF mapping revealed the representation of the lower right visual quadrant, from the fovea out to about six degrees of visual angle (18, 19). The employed grating patch resulted in a topographic map of visually induced gamma-band power with a clear peak at the V1 representation of the stimulus (Fig. 2A).

**Fig. 2.**
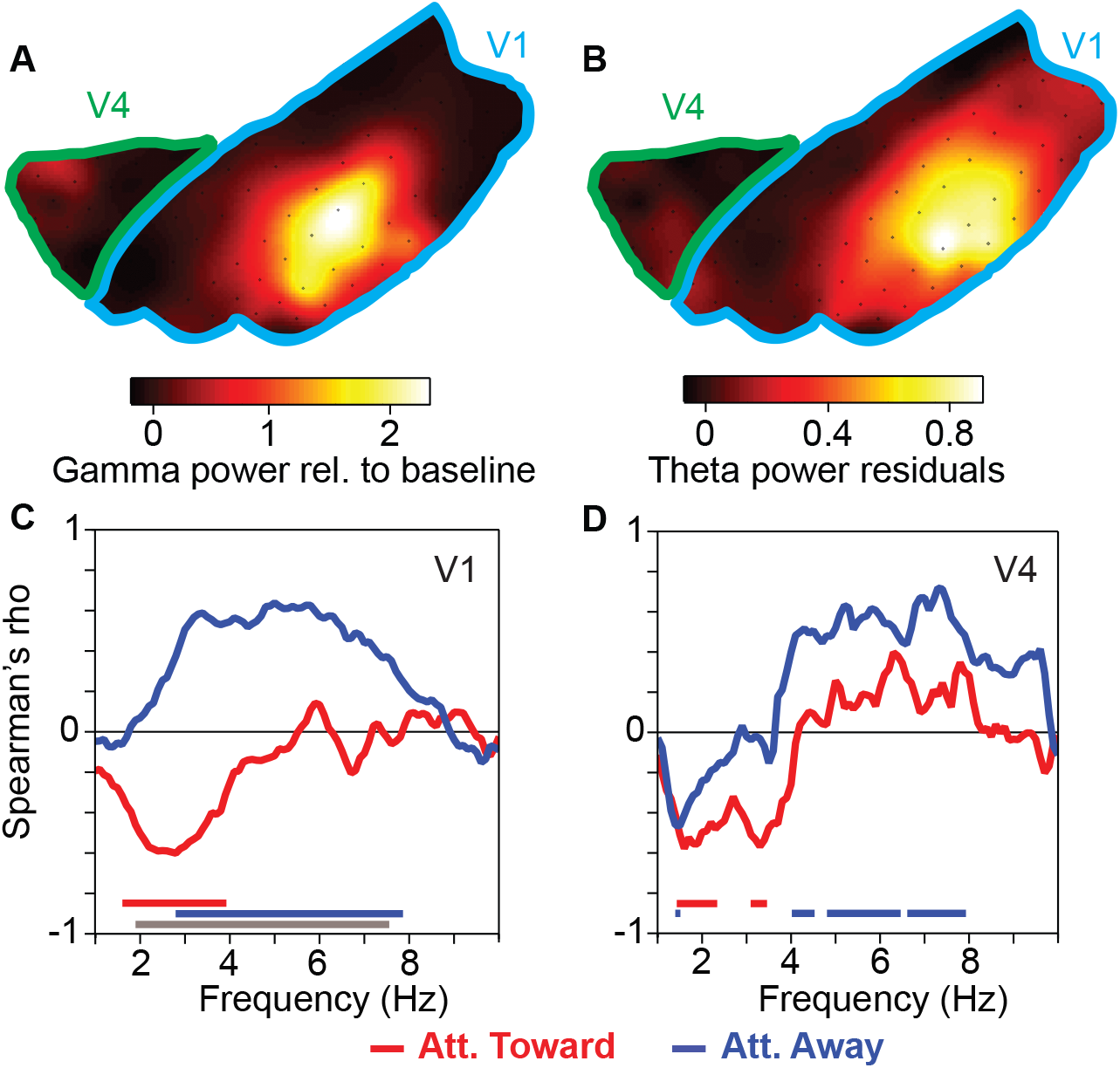
The theta rhythm coextends with the visually induced gamma rhythm. (*A*) Visually induced LFP gamma-band power, as a function of spatial location in V1-V2 (indicated by blue outline) and V4-TEO (indicated by green outline). (*B*) Same as *A*, but showing LFP theta-band power after removing the 1/f^n^ component. (*C*) Correlation between 1) visually induced gamma-band power and 2) the power (1/f^n^ removed) at the frequency indicated on the x-axis, across recording sites in V1-V2. Colored lines on the bottom indicate frequencies with significant correlations with attention toward (red) or away from (blue) the activating stimulus (p<0.05; non-parametric permutation test with correction for multiple comparisons across frequencies). The gray line on the bottom indicates frequencies with a significant difference in correlation between the attention conditions (same test). (*D*) Same as *C*, but for area V4.

When we calculated a corresponding topographic map of theta power, it also showed a clear spatial peak (Fig. 2B). For this analysis, the 1/f^n^ component was estimated by robust regression and it was subtracted (20). For the theta power analysis, the 1/f^n^ component constitutes noise, and its subtraction attenuates noise differences between electrodes and monkeys and thereby improves the signal-to-noise ratio for the theta component. Also, we used the trials, in which the stimulus represented by the recorded part of V1 was non-attended, because these “non-attended trials” contained stronger theta, as we will show below. Theta power and visually induced gamma power were spatially coextensive. We calculated the spatial correlation (Spearman rank correlation) between theta power (1/f^n^ component removed) and visually induced gamma power, across electrodes, for each of the low frequency components up to 10 Hz. The resulting correlation spectra reveal that across the post-lunate ECoG, visually induced gamma is positively correlated with theta, when the stimulus is non-attended (Fig. 2C, blue line). Anterior to the lunate, between lunate and superior temporal sulcus (STS), the ECoG covered superficial V4 and parts of area TEO. Even though retinotopy in this region is coarser, the spatial correlation spectrum was similar, when the stimulus was non-attended (Fig. 2D, blue line). When the stimulus was attended, gamma was negatively correlated with power around 1-4 Hz, in both the post- and pre-lunate region (Fig. 2C,D, red lines). To ensure that the correlations shown in Figure 2C, D are not due to broadband power correlations, the analyses used gamma from the pre-cue period and theta power from the post-cue period (both with visual stimulation), that is, from non-overlapping trial epochs. Results are essentially the same if the post-cue period is used for both.

For all further analyses, we selected the electrodes that were most strongly visually driven, concretely, we selected the 25% of electrodes with the strongest visually induced gamma-band activity, separately for the post- and the pre-lunate region. In the post-lunate region, all selected electrodes were likely on V1, and we refer to them simply as V1 (without each time referring to the fact that only the selected electrodes were used). In the pre-lunate region, selected electrodes were on V4, maybe including posterior TEO, yet for simplicity, we will refer to them as V4 (again implying the selection). For the respective phase-locking analyses, we used all sites (bipolar derivations), that included a selected electrode.

### Selective attention reduces theta

Raw power spectra averaged over V1 electrodes showed a peak around 4 Hz (Fig. 3A). This V1 theta rhythm was reduced when the visual stimulus was attended. A similar pattern was found in V4: There was a shallow bump close to 4 Hz, which was reduced by attention (Fig. 2B). To reveal the peaks and their peak frequencies more clearly, we again estimated and removed the 1/f^n^ component by robust regression (20). The residual power spectra showed distinct peaks around 4-5 Hz in both V1 and V4, which were strongly reduced by attention (Fig. 3C,D). The theta rhythms in V1 and V4 can also be seen in a time-frequency analysis (Fig. S1).

**Fig. 3.**
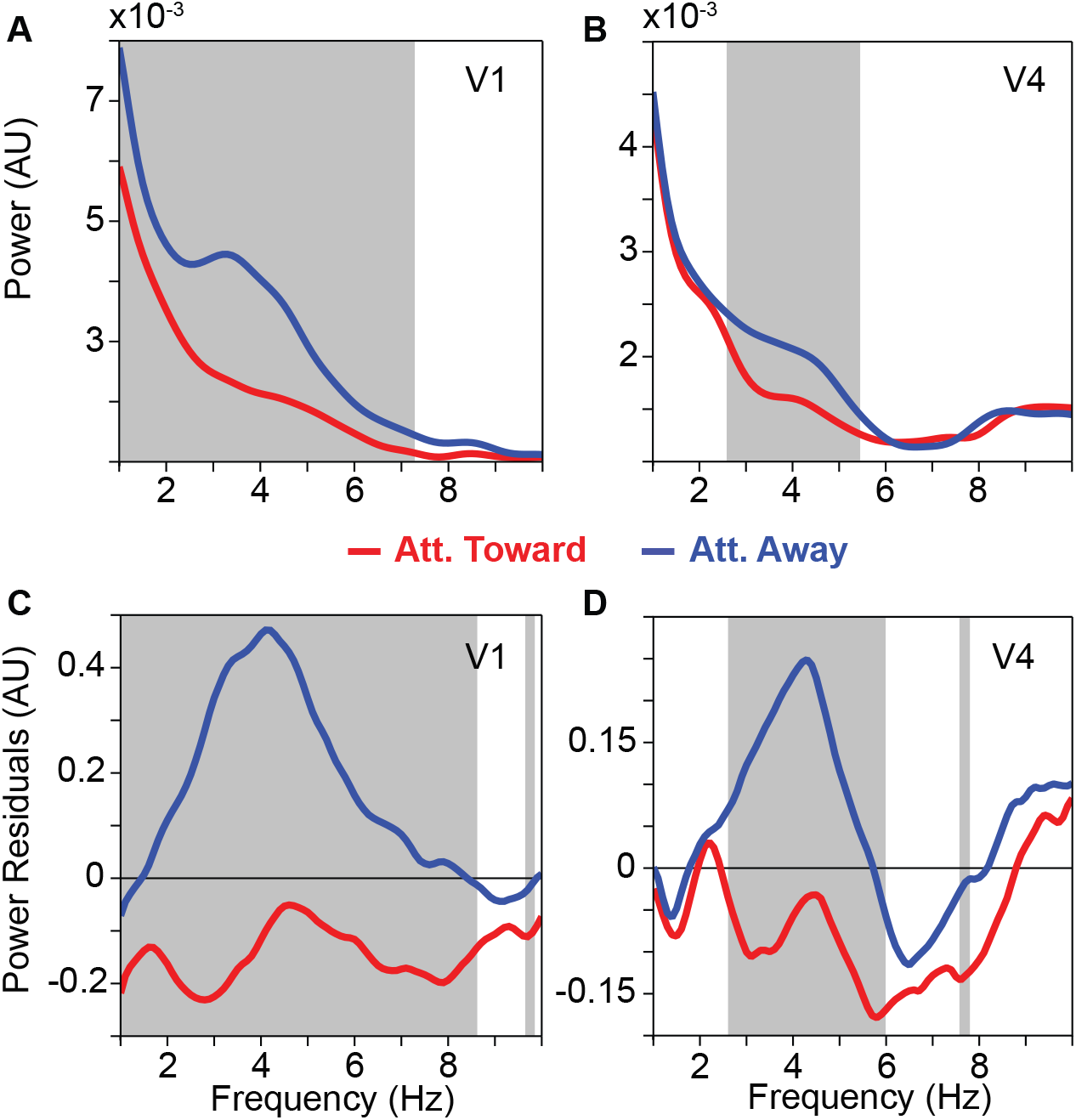
Average low-frequency LFP power spectra and their modulation by selective attention. (*A*) Average LFP power spectra in area V1 with attention toward (red) and away (blue) from the activating stimulus. The gray-shaded region indicates frequencies with a significant difference between attention conditions (p<0.05; non-parametric permutation test with correction for multiple comparisons across frequencies). (*B*) Same as *A*, but for area V4. (*C*) Same as *A*, but showing the power residuals after removing the 1/f^n^ component of the power spectrum through robust regression (see Materials and Methods). (*D*) Same as *C*, but for area V4.

Note that the increased theta power in the attend-away condition most likely reflects an increased theta-rhythmic synchronization among local neurons. In general, power increases can be due to increases in synchronization or increases in the firing rates of the involved neurons. However, V1 and V4 neurons, when activated with one stimulus in their RFs as done here, show either increases or no change in their firing rates with attention (21). Thus, the enhanced theta power in the attend-away condition cannot be due to enhanced neuronal firing rates.

Phase-locking (PPC) spectra for pairs of sites within V1 or V4 and between V1 and V4 showed corresponding theta peaks. When the stimulus was attended, this theta-phase locking was strongly reduced but not fully eliminated (Fig. 4).

**Fig. 4.**
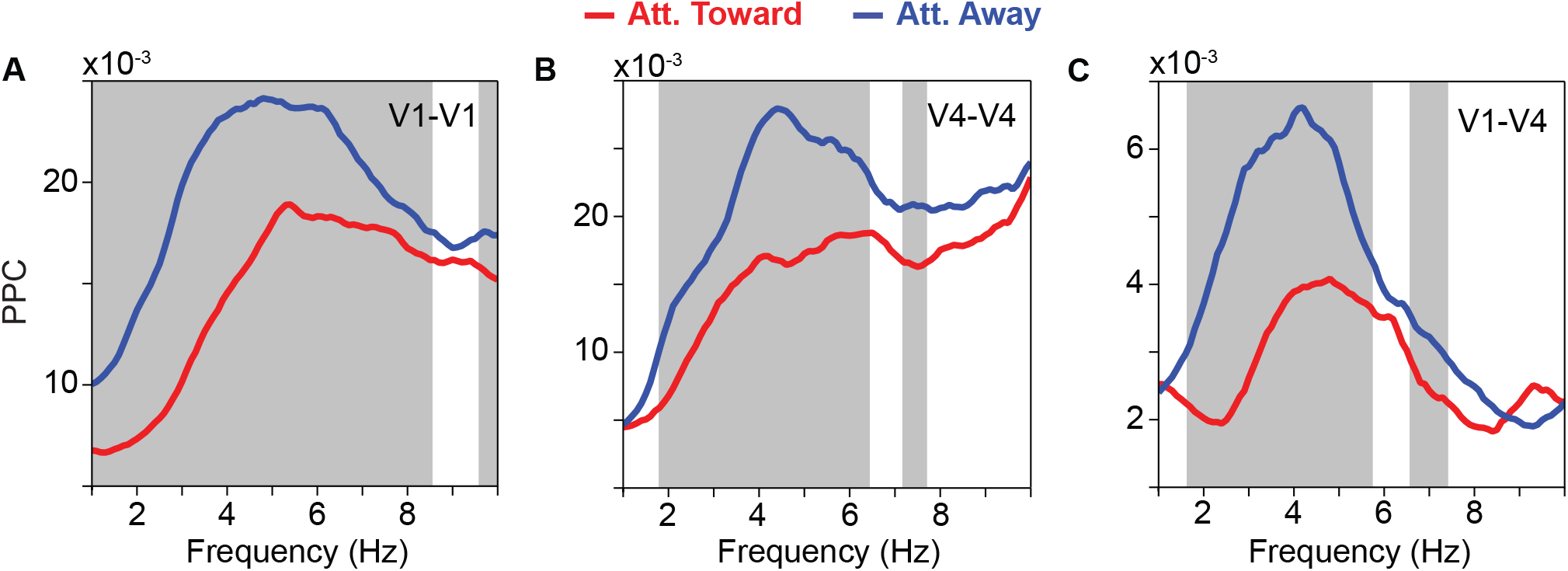
Average low-frequency LFP phase-locking (PPC) spectra and their modulation by selective attention. (*A*) Average LFP phase locking between sites within area V1 with attention toward (red) and away from (blue) the activating stimulus. The gray-shaded region indicates frequencies with a significant difference between attention conditions (p<0.05; non-parametric permutation test with correction for multiple comparisons across frequencies). (*B*) Same as *A*, but between sites within area V4. (*C*) Same as *A*, but between sites in area V1 and sites in area V4.

### Theta-band Granger causality is stronger in the feedforward direction and reduced by attention

We next investigated Granger causality (GC) between V1 and V4 in the low-frequency range. Figure 5A shows the GC spectra averaged over all interareal pairs of sites between V1 and V4, pooled across both attention conditions, separately for the feedforward (V1-to-V4; green line) and feedback (V4-to-V1; black line) direction. These GC spectra reveal clear theta peaks, and show that the theta GC is stronger in the feedforward than feedback direction.

The GC metric can be influenced by noise in the two signals. Such an influence can be diagnosed by reversing the time axis, because this keeps the signals (with their contained noise) identical, while reversing the temporal relation (22). Thus, an asymmetry in GC that is genuine should invert upon time reversal. We performed this test on our data, and the asymmetry in GC between V1 and V4 did indeed invert, suggesting that the asymmetry is genuine (Fig. S2A).

**Fig. 5.**
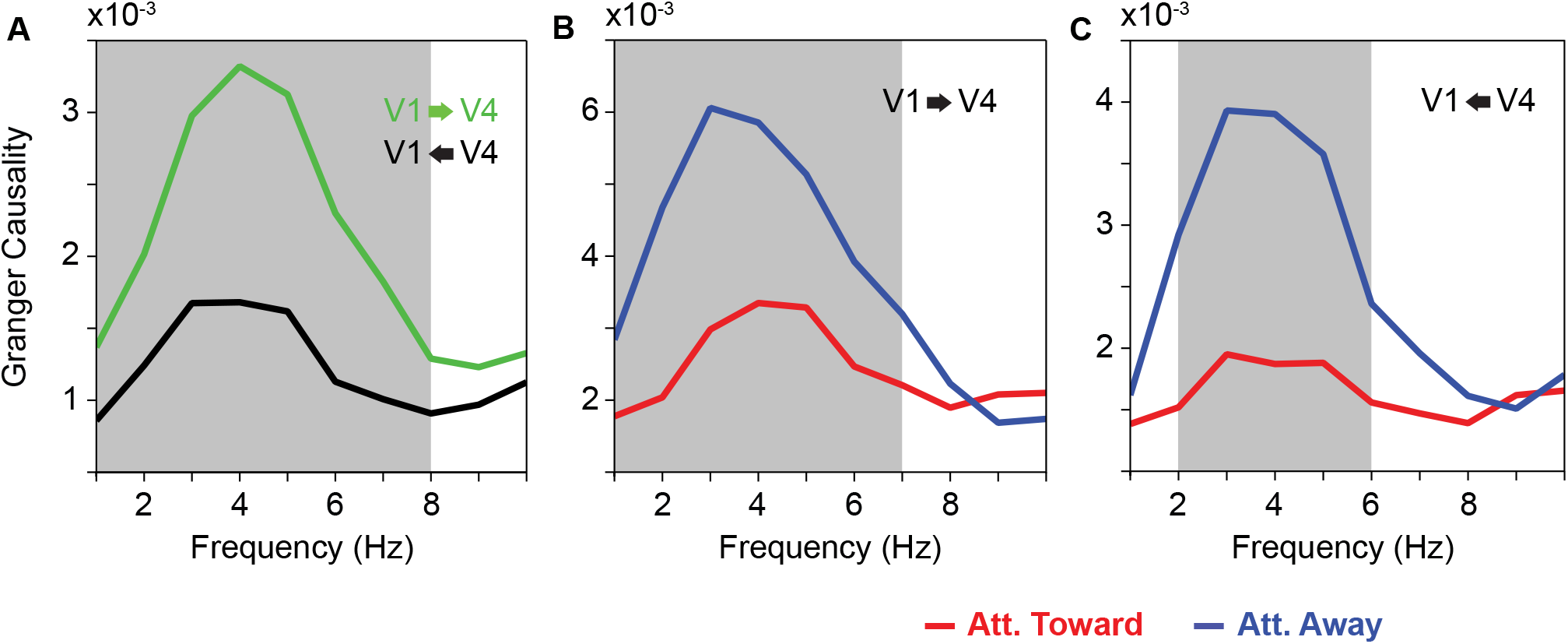
Average low-frequency Granger causality (GC) spectra between V1 and V4 sites. (*A*) Average GC-influence spectra between V1 and V4 in the feedforward (green) and feedback directions (black). The gray-shaded regions indicate frequencies with a significant difference between bottom-up and top-down (p<0.05; non-parametric permutation test with correction for multiple comparisons across frequencies). (*B*) Average GC-influence spectra between V1 and V4 in the feedforward direction, with attention toward (red) and away from (blue) the activating stimulus. The gray-shaded regions indicate frequencies with a significant difference between attention conditions (p<0.05; non-parametric permutation test with correction for multiple comparisons across frequencies). (*C*) Same as *B*, but for the feedback direction.

Figure 5B,C shows the GC spectra separately for the two attention conditions. When the stimulus was attended, both feedforward and feedback GC were reduced in the theta band.

### Theta-gamma phase-amplitude coupling and its attentional modulation

Several previous studies in other brain areas have reported gamma power to be modulated with theta phase, that is, they reported theta-gamma phase-amplitude coupling, or PAC (23–26). We investigated whether the theta rhythm that we found in V1 and V4 modulates gamma power, and whether this is affected by selective attention. Figure 6A shows for one example electrode location, and for attention away from the stimulus, the raw spectral power as a function of time relative to the theta trough. This reveals that the amplitude of visually induced gamma-band power is modulated systematically by theta phase. Figure 6B shows the resulting PAC, averaged over V1 and V4, and over both attentional conditions. It reveals a distinct PAC peak between theta phase and gamma power. Note that the theta-rhythmic modulation of gamma was most pronounced for the high-frequency end of the gamma band. In addition, this analysis revealed PAC between the phase around 1 Hz and power in several frequency bands; this 1 Hz component is likely related to the temporal frequency of the drifting gratings (see Materials and Methods). Note further that gamma amplitude was also modulated with the phase in the alpha band (10-12 Hz).

**Fig. 6.**
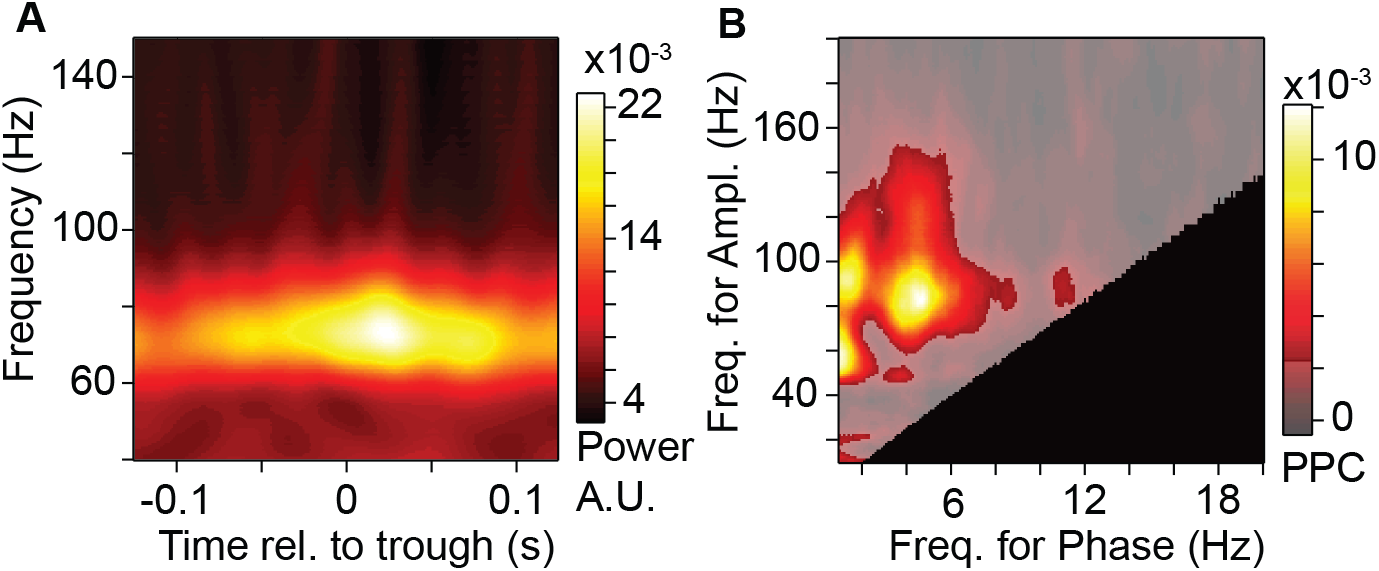
Theta-gamma phase-amplitude coupling (PAC) in visual cortex. (*A*) LFP power of one example site in the 40-150 Hz range (y-axis) as a function of time relative to the theta trough (x-axis). (*B*) Grand-average PAC as a function of the frequency defining the power (y-axis) and the frequency defining the phase (x-axis). The semitransparent gray mask indicates frequency pairs with non-significant PAC (p<0.05; non-parametric permutation test with correction for multiple comparisons across frequency pairs). The black area indicates frequency pairs excluded from the analysis (see Materials and Methods).

Area V1 showed a distinct theta-gamma PAC peak, which was strongly reduced by attention (Fig. 7A-C). At the same theta-gamma phase-amplitude frequencies, area V4 showed significant PAC when attention was away from the stimulus, yet no significant PAC difference between attention conditions (Fig. 7D-F). Also, V4 gamma power was modulated with V1 theta phase, when attention was away from the stimulus, but not when attention was to the stimulus; yet this attentional contrast did not reach significance (Fig. 7G-I). There was no significant modulation of V1 gamma power by V4 theta phase. Additional PAC components occurred at lower phase frequencies and were likely related to the temporal frequency of the drifting gratings, as mentioned above.

**Fig. 7.**
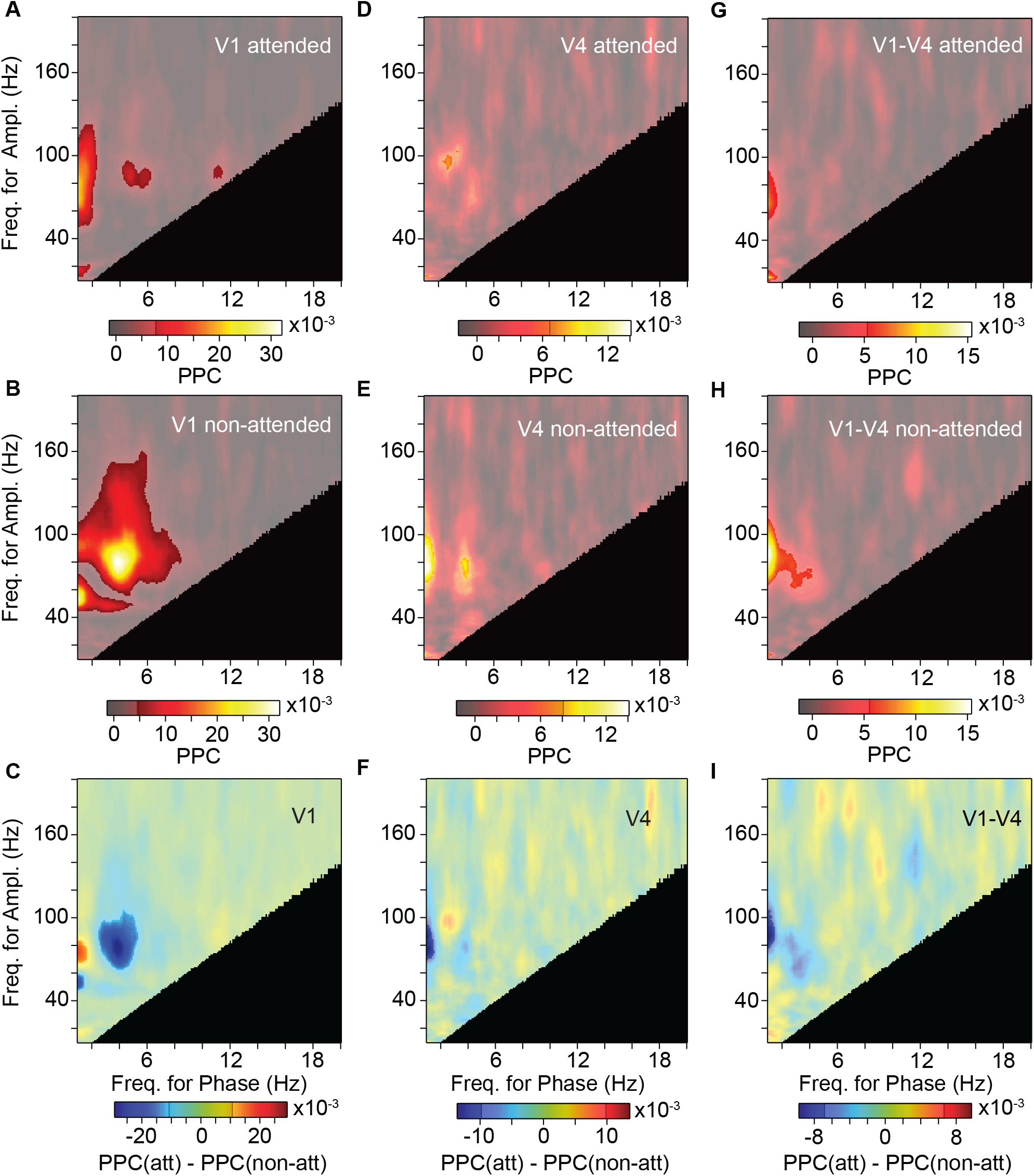
Modulation of PAC by selective attention. (*A, B*) Average PAC in area V1 with attention toward (*A*) and away from (*B*) the activating stimulus. (*C*) Average PAC difference in area V1 between the two attention conditions shown in *A* and *B*. The semitransparent gray mask indicates frequency pairs with non-significant PAC, or PAC difference, respectively (p<0.05; non-parametric permutation test with correction for multiple comparisons across frequency pairs). The black area indicates frequency pairs excluded from the analysis (see Materials and Methods). (*D*), (*E*), (*F*), Same as *A, B, C*, but for area V4. (*G*), (*H*), (*I*), Same as *A, B, C*, but using the phase in area V1 and the amplitude in area V4.

The quantification of PAC entails the estimation of time-varying power. Power estimation with spectral specificity requires windows of some length. This leads to low-pass filtering of power time courses. For the main analyses, we eliminated phase-amplitude pairs, for which the low-pass filtering reduced power to less than 70% of the power in the passband (black regions in PAC plots; see Methods for details). When we lowered the threshold to 30%, this allowed to investigate more phase-amplitude pairs, but it did not reveal additional significant PAC, in particular no theta-beta PAC.

### Visual theta remains after microsaccade removal

It has previously been shown that theta-band rhythmicity is present in the sequence of microsaccades (MSs) (27, 28). MSs cause a movement of the retinal image and an MS-related response in the LFP and the multi-unit activity (27). MSs also modulate the strength of gamma-band activity (27, 28). Thus, the MS rhythm may underlie both the theta rhythm and the theta-gamma PAC observed here. To investigate this, we first quantified the phase-locking between MSs and the LFP in V1. Figure 8A shows the MS-LFP PPC spectrum and reveals a clear theta peak. If neuronal activity and phase locking in the theta band were due to driving by theta-rhythmic MSs, then removal of epochs with MSs should diminish the observed theta rhythmicity. To test this, we excluded MSs and investigated the effect on the observed neuronal theta rhythmicity. We detected MSs and excluded data recorded between MS onset and 0.5 s thereafter. This substantially reduced the amount of available data. We calculated low-frequency PPC spectra within V1 for the attend-away condition for 1) all available epochs (N=1917 epochs), 2) epochs excluding MSs exceeding average eye speed by 5 SD (N=827 epochs). Figure 8B reveals that excluding MSs did not decrease theta rhythmicity in V1 (note that the PPC metric has no sample-size bias (17)). This result strongly suggests that, while there is phase locking between MSs and visual cortical theta, the theta exists independently of the occurrence of MSs.

**Fig. 8.**
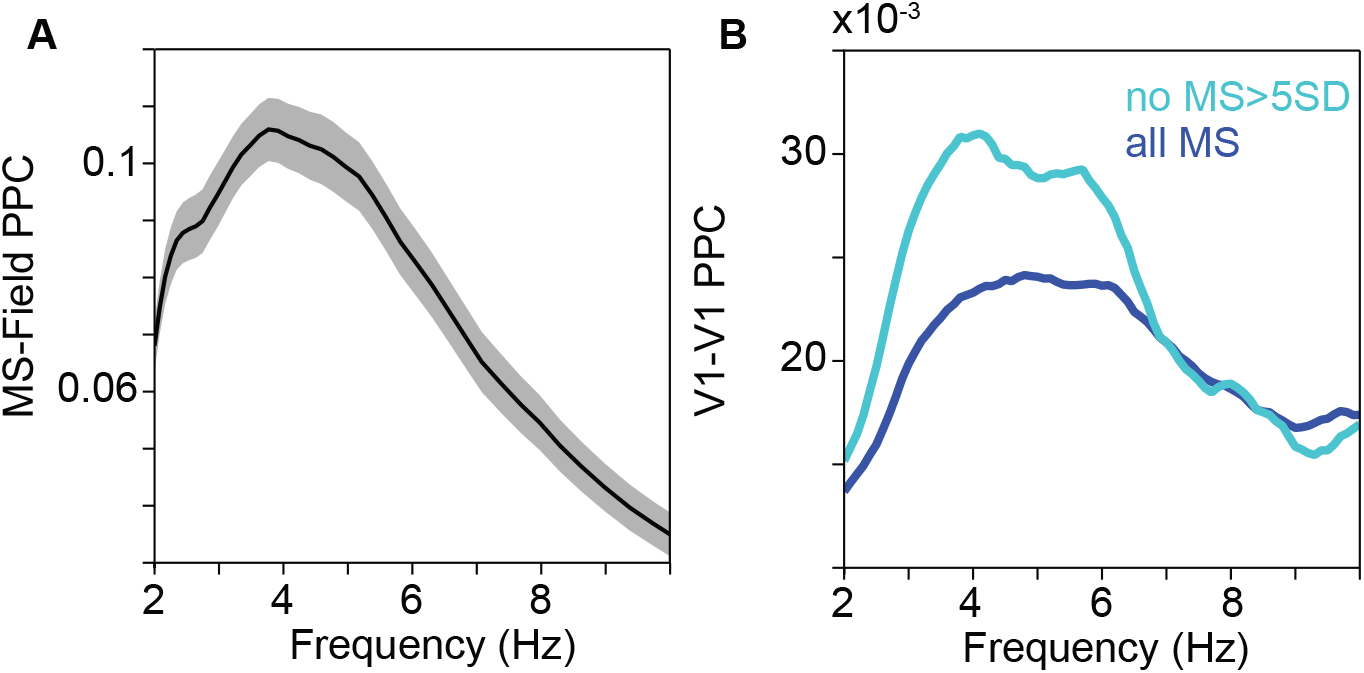
Visual theta remains after microsaccade removal. (*A*) MS-LFP PPC as a function of frequency, showing a clear theta peak. The shading shows the 95% confidence interval based on a bootstrap procedure across MSs. (*B*) V1-V1 PPC in the attend-away condition, as a sensitive metric of V1 theta rhythmicity, calculated after removing epochs with MSs. Dark blue line: All available epochs (N=1917); Light blue line: Epochs excluding MSs, that exceeded the mean eye speed by 5 SD (N=827).

Figure 9 investigates the influence of this MS removal on further metrics of visual theta. The main results remained essentially unchanged. Power spectra (with and without robust regression and removal of the 1/f^n^ component) showed theta peaks when attention was away from the stimulus. Those theta peaks were strongly reduced by attention to the stimulus (Fig. 9A-D). PPC spectra showed theta peaks, and those peaks showed attentional reductions that reached significance for V1-V1 and V1-V4 and tended in the same direction for V4-V4 (Fig. 9E-G). GC spectra confirmed the predominant feedforward influence (Fig. 9H). PAC in V1 showed a theta-gamma peak, which was significantly reduced by attention (Fig. 9G,H,I). PAC in V4 and between V1 phase and V4 power lost significance, both per condition and in the condition difference, probably due to reduced sensitivity (Fig. S3).

**Fig. 9.**
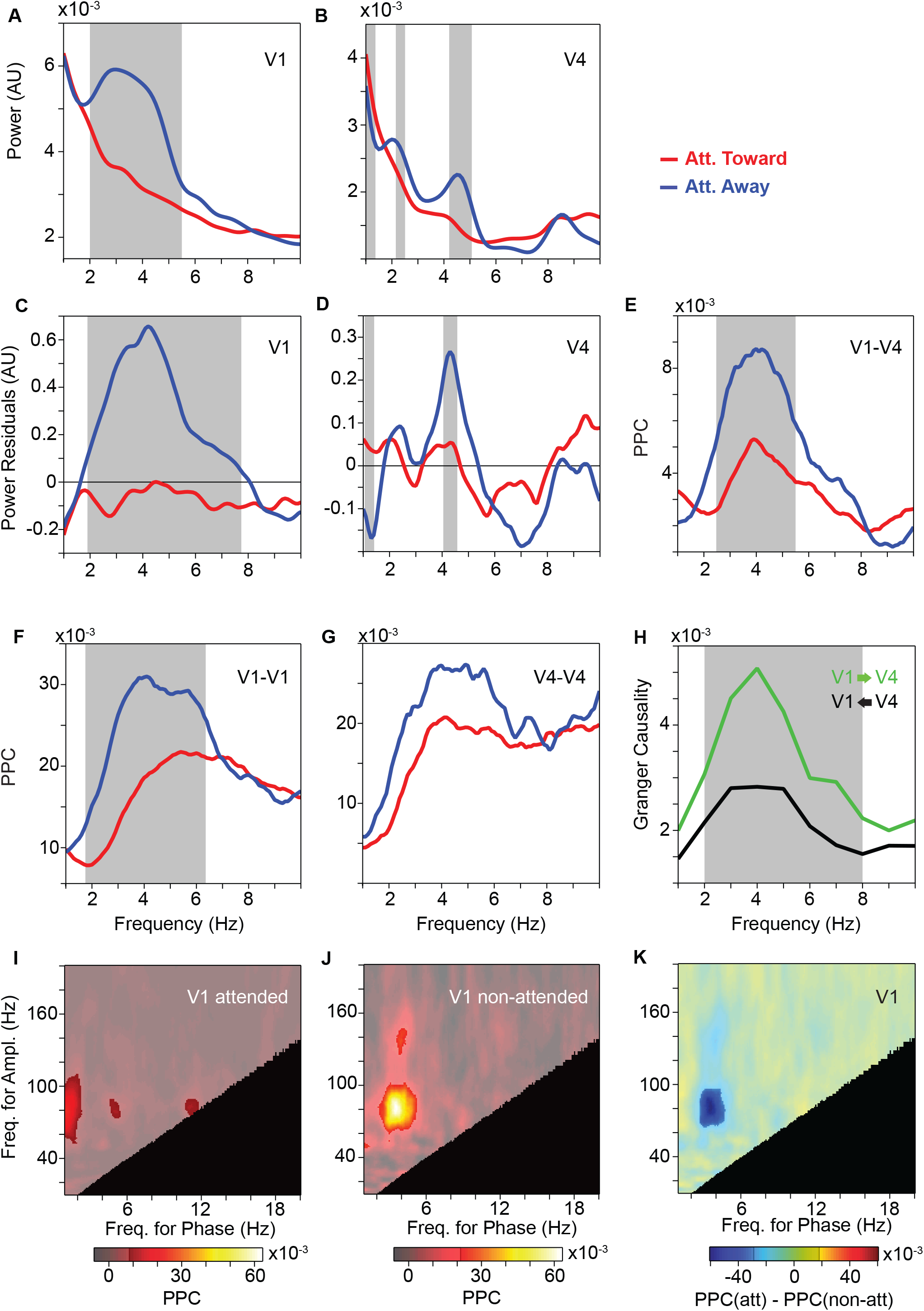
Attention contrast, excluding epochs with microsaccades. (*A*) Average LFP power spectra in area V1, with attention toward (red) and away (blue) from the activating stimulus. (*B*) Same as *A*, but for area V4. (*C*), (*D*) Same as *A* and *B*, after removing the 1/f^n^ component. (E) Average LFP phase locking between sites in area V1 and sites in area V4, with attention toward (red) and away from (blue) the activating stimulus. (*F*), (*G*) Same as *E*, but for pairs of sites within area V1 (*F*) and area V4 (*G*). (*H*) Average GC-influence spectra between V1 and V4 in the feedforward (green) and feedback directions (black). (*A-H*) The gray-shaded region indicates frequencies with a significant difference between attention conditions (*A-G*), or between feedforward and feedback directions (*H*) (p<0.05; non-parametric permutation test with correction for multiple comparisons across frequencies). (*I*), (*J*) Average PAC in area V1 with attention toward (*I*) and away from (*J*) the activating stimulus. (K) Average PAC difference in area V1 between the two attention conditions. (*I-K*) The semitransparent gray mask indicates frequency pairs with non-significant PAC, or PAC difference, respectively (p<0.05; non-parametric permutation test with correction for multiple comparisons across frequency pairs). The black area indicates frequency pairs excluded from the analysis (see Materials and Methods).

### Control for microsaccade rate

In addition, we performed an alternative control, by equating the MS rate, that is, the MS temporal density, between attention conditions. Figure S4A shows the cumulative distribution of MS rate over the respective number of data epochs. MS rate actually differed between attention conditions. We therefore stratified the data to arrive at two equally sized sets of epochs with an essentially equal distribution of MS rates (dashed lines in Fig. S4A). After stratification, almost all main results remained essentially unchanged (Fig. S4B-K).

### Control for theta power

Finally, we controlled for the possibility that the effects of attention on theta PPC, theta GC or theta-gamma PAC were explained by the effects of attention on theta power. Theta power was reduced by attention, which might reduce the sensitivity of PPC, GC and/or PAC quantification, which might in turn explain the reduced PPC, GC and/or PAC values with attention. To investigate this possibility, we stratified for theta power between attention conditions. After stratification, attention conditions still differred significantly in V1 theta-gamma PAC (Fig. S5A), but they did not any more differ significantly in theta PPC (V1-V1, V4-V4 or V1-V4) (Fig. S6), theta GC (V1-to-V4 or V4-to-V1) (Fig. S2B,C), theta-gamma PAC in V4 (Fig. S5B), or theta-gamma PAC between V1 phase and V4 gamma (Fig. S5C). Where the stratification rendered the attention contrast non-significant, this is consistent with two interpretations. One interpretation relates to the signal-to-noise ratio (SNR) of theta. The enhanced theta power in the attend-away condition might increase the sensitivity of the theta PPC and the theta-gamma PAC quantification and thereby explain the attention effect on those metrics. An alternative interpretation relates to the relative amount of time with strong theta power. The enhanced theta power in the attend-away condition might correspond to more time spent in a regime of strong theta rhythmicity. This might conceivably be a genuine difference between attention conditions. If this is the case, stratification for theta power artificially removes this genuine difference. There is no unequivocal way to distinguish between these two interpretations.

## Discussion

We demonstrate the presence of a ≈4 Hz theta rhythm in awake macaque visual cortex. This theta rhythm was present selectively in sites driven by the visual stimulus, such that the spatial map of theta was correlated with the map of visually induced gamma-band activity. Theta rhythmically modulated local gamma-band activity and thereby most likely the gamma-associated local processing of visual information. Theta rhythms in V1 and V4 synchronized, and an analysis of GC revealed a predominant feedforward influence. Theta rhythmicity was substantially reduced when the visual stimulus was attended. Visual cortical theta showed phase locking with MSs. Yet, exclusion of MS effects left all main theta-related observations essentially unchanged.

We were somewhat surprised to find that theta shows a clear spatial correlation or coextension with visually induced gamma-band power. There were reasons to assume that a putative theta rhythm might be global across visual cortex. Hippocampal recordings suggest that theta is global in this structure, travelling as a wave from dorsal to ventral parts (29, 30). Also, there is the general notion that slower rhythms are more global than faster rhythms (31, 32). Yet, we find theta to be spatially specific, coupled to gamma by spatial extension and also through PAC. This link between theta and gamma is reminiscent of the finding that inter-areal GC influences in both theta and gamma are typically stronger in the anatomically defined feedforward than feedback direction (33).

The PAC analysis showed theta-gamma coupling that peaked for an amplitude-frequency at the high-frequency end of the visually induced gamma band activity. Thus, theta-rhythmic modulation was most apparent for this high-frequency part of the overall gamma peak. This might reflect a physiological asymmetry or be related to signal-to-noise ratios. Physiologically, it is conceivable that the modulation is in fact stronger at the upper flank of the gamma peak than at the lower flank, which would be equivalent to an asymmetric broadening of the gamma peak towards higher frequencies. Alternatively, the gamma-band peak is modulated in its entirety, yet the PAC metric ends up larger for the upper than the lower flank, e.g. because the gamma peak is superimposed on unmodulated (or less modulated) 1/f^n^ power. If we consider the 1/f^n^ component of the power spectrum as noise, this noise is larger for the lower than the upper flank.

In addition, it is interesting to investigate the precise frequency of the observed theta rhythm. The basic spectra of power (residuals) and phase locking showed peaks close to 4 Hz. The analysis of spatial correlation between theta power and visually-induced gamma power showed a broader peak that includes 4 Hz, yet extends up to 8 Hz. This suggests that the underlying phenomenon might actually occupy this broader frequency range, with theta merely peaking at 4 Hz for the particular stimulus and task conditions used here. Whether other stimuli or tasks make theta in V1 and/or V4 shift in frequency is an interesting topic for further study. In any case, the 4-8 Hz range found in the spatial correlation analysis is an interesting link to the classical hippocampal theta, which occupies this range. Hippocampal theta in fact shifts in frequency, e.g. depending on running speed (34, 35).

The mechanisms behind the observed visual cortical theta rhythm and its attentional modulation are not known. Theta and its mechanisms have probably been most intensively studied in medial temporal lobe (MTL), in particular the hippocampus and entorhinal cortex (2, 3, 29, 30, 36–39). Hippocampal theta is partly synchronized to neocortex, e.g. to entorhinal and prefrontal areas (23, 40, 41). Theta in PFC further synchronizes with strongly connected structures like the anterior cingulate cortex (ACC) and the posterior parietal cortex (42, 43). The theta rhythm in non-human primate PFC shows long-distance synchronization to a theta rhythm in area V4 (44). In macaque area MT, the power of high-frequency (30-120 Hz) LFP components is modulated by the phase of low-frequency (1-8 Hz) components, and this modulation is reduced by attention (45). Thus, in principle, the theta described here could originate in hippocampus and progress via PFC and extrastriate visual areas to V1. Yet, this seems implausible, because such a mechanism would most likely not generate the observed spatial coextension between theta and gamma, and the predominant GC direction from V1 to V4. Moreover, the present results place further constraints on potential mechanisms: The fact, that removing MSs left the main results essentially unchanged, suggests that theta in visual cortex does not merely reflect theta-rhythmic MSs. Rather, the clear spatial co-extension between theta power and visually induced gamma suggests a role for visually driven activity in theta generation.

The theta rhythms in V1 and V4 were reduced by selective attention. Attention effects are typically smaller in V1 than in higher visual areas (for otherwise comparable conditions). This holds for firing rates (21, 46) and gamma-band synchronization (47). In fact, for gamma-band synchronization, different studies in V1 have reported attentional increases (47), decreases (48) or the absence of an effect (18). By contrast, the attentional effects on theta appeared to be of similar strength in V4 and V1, and thereby establish an unusually strong attention effect for V1.

Many studies have reported reductions in alpha power at the neuronal representation of visual stimuli or visual attention (49, 50). The attentional reduction of theta observed here might appear like a related phenomenon at a slightly lower frequency. However, whereas visually driven neuronal ensembles show reduced alpha (50), we found that they show enhanced theta (Fig. 2). This observation supports an alternative scenario. Recent studies have shown that attention samples visual stimuli at a theta rhythm. When human subjects have to detect the appearance of a faint stimulus at a peripheral location, their detection performance is modulated by the phase of a 7-8 Hz rhythm with a maximum over frontal cortex (51). This might reflect an ≈8 Hz rhythmic attentional sampling. In support of this, three subsequent studies have shown that two simultaneously monitored stimuli are attentionally sampled in alternation, each at ≈4 Hz (8, 11, 12). A further study estimated the temporal sampling frequency of attention, and found it to be around 7 Hz for a single attended stimulus, 4 Hz for two and 2.6 Hz for three (52). These numbers are consistent with a single attentional sampling mechanism at ≈8 Hz that is multiplexed over the to-be-attended stimuli. Such a scenario would also explain theta-rhythmic modulations of firing rates in inferotemporal (IT) cortex during the presentation of two stimuli (14). When IT neurons respond to one stimulus, and a second stimulus is added onto the screen, firing rates start oscillating at ≈4-6 Hz in a way that suggests that attention is drawn to the newly presented stimulus and subsequently alternates between the two stimuli.

At first glance, these results might seem to suggest that visual cortical theta should be stronger for the attended stimulus. However, the fact that divided attention tasks reveal theta-rhythmic sampling, does not mean that attended stimuli are affected by stronger theta-rhythmic modulation than non-attended stimuli. The evidence so far suggests different effects of a frontal ≈7-8 Hz rhythm and a visual cortical ≈4-6 Hz rhythm. The original study, reporting a modulation of visual stimulus detection by the frontal ≈7-8 Hz EEG phase, found this effect to be present for attended but not for unattended stimuli (51). By contrast, a recent study using a very similar approach, but focusing on the ≈4-6 Hz EEG phase in occipital electrodes contralateral to the respective stimulus, with retinotopically selective responses, revealed a stronger effect for unattended than attended stimuli (53). Another recent study used two independent white-noise sequences as stimuli to calculate their respective temporal response functions (54). This revealed that sampling is more prominent when attention is distributed to two stimuli as compared to when it is focused on one stimulus. These recent results from human subjects are consistent with the abovementioned recordings in macaque IT cortex, which showed a ≈4-6 Hz rhythm to be strong when two stimuli are presented, and weaker when a single stimulus is presented and thereby fully attended (14). Thus, a number of studies, including the present one, suggest that attentional processes in visual cortex are mainly sustained, yet still weakly theta rhythmic, at the attended location, and that attention theta-rhythmically scans the remaining visual space to explore other stimuli. As a consequence, non-attended stimuli receive attentional processing benefits only when they are attentionally scanned, leading to relatively strong theta rhythmicity.

Future studies will need to investigate whether attentional control structures show an ≈7-8 Hz sampling rhythm that is coherent to the sampled stimulus representations in visual cortex. As mentioned above, the ≈7-8 Hz EEG component, whose phase predicts human detection performance, is strongest over frontal areas (51). Also, spike and LFP recordings in macaque parietal cortex have recently revealed a similar theta rhythm (42, 55). If such theta-rhythmic top-down influences were to be found, it will be interesting to understand how they fit with the predominantly bottom-up directed theta influences observed between visual areas (33). One possibility is that control structures exert a theta-rhythmic perturbation on early and even primary visual cortex, which then percolates up through the hierarchy of visual areas.

## Methods

### Subjects, stimuli and task

Two adult male macaque monkeys participated in this study. All procedures were in accordance with Dutch and European regulations for the protection of animals and were approved by the animal ethics committee of Radboud University Nijmegen (Netherlands). The data analyzed here have been (partially) used in previous studies (15, 18, 19, 22, 33, 56–61).

Stimuli and behavior were controlled by the software CORTEX (http://dally.nimh.nih.gov). Stimuli were presented on a CRT monitor at 120 Hz non-interlaced. When the monkey touched a bar, a gray fixation point appeared at the center of the screen. When the monkey brought its gaze into a fixation window around the fixation point (0.85 degree radius in monkey K; 1 deg radius in monkey P), a pre-stimulus baseline of 0.8 s started. If the monkey’s gaze left the fixation window at any time, the trial was terminated. The measured eye positions during correct trials used for analysis differed only by an average of 0.03 deg of visual angle between the two attention conditions. After the baseline period, two physically isoluminant patches of drifting sinusoidal grating appeared (diameter= 3 degrees, spatial frequency ≈1 cycle/degree, drift velocity ≈1 degree/s, resulting temporal frequency ≈1 cycle/s, contrast = 100%). The two grating patches chosen for a given recording session always had equal eccentricity, size, contrast, spatial frequency and drift velocity. The two gratings always had orientations that were orthogonal to each other, and they had drift directions that were incompatible with a Chevron pattern moving behind two apertures, to avoid pre-attentive binding. In any given trial, one grating was tinted yellow, the other blue, with the color assigned randomly across trials. The yellow and blue colors were physically equiluminant. After 1-1.5 s (0.8-1.3 s in monkey P), the fixation point changed color to match the color of one of the two gratings, thereby indicating this grating as the relevant stimulus and the other as irrelevant. For each trial, two independent change times for the two stimuli were determined randomly between stimulus onset and 4.5 s after cue onset, according to a slowly rising hazard rate. If the relevant stimulus changed (before or after the irrelevant stimulus changed), and the monkey released the bar within 0.15-0.5 s thereafter, the trial was terminated and a reward was given. If the monkey released the bar at any other time, the trial was terminated without reward. The stimulus changes were small changes in the grating pattern, with the stripes undergoing a gentle bend. During the bend, the outer ends of the grating stripes lagged increasingly behind the center of the stripes, until the lag reached 0.1 degree at 75 ms after the start of the bend. Over the course of another 75 ms, the stripes straightened again.

Several sessions (either separate or after attention-task sessions) were devoted to the mapping of receptive fields (RFs), using 60 patches of moving grating. Receptive field positions were stable across recording sessions (18).

### Neurophysiological recordings and signal preprocessing

Neuronal recordings were made from two left hemispheres in two monkeys through a micromachined 252-channel electrocorticographic electrode array (ECoG) implanted subdurally. The details of the production and the electrochemical properties have been described in a separate paper (62). Briefly, ECoG grids were 10 micron thick polyimide foils with 0.3 micron thick Platinum electrodes and conductive lanes embedded. Electrodes had an exposed surface with a diameter of 1 mm and a center-to-center spacing of 2-3 mm. Electrodes were arranged in lanes, and two neighboring lanes ran parallel on one “finger” of the polyimide foil (33). The structuring in separate fingers avoided wrinkling of the ECoG on the brain surface and corresponding pressure points. For ECoG implantation, a 6.5×4 cm craniotomy over the left hemisphere in each monkey was performed under aseptic conditions with isoflurane anesthesia. The dura was opened and the ECoG was placed directly onto the brain under visual control. Several high resolution photos were taken before and after placement of the ECoG for later coregistration of ECoG signals with brain regions. After ECoG implantation, both the dura and the bone were placed back and secured in place. After a recovery period of approximately three weeks, we started with neuronal recordings.

Signals obtained from the 252-electrode grid were amplified 20 times by eight Plexon headstage amplifiers (Plexon, USA), high-pass filtered at 0.159 Hz, low-pass filtered at 8 kHz and digitized at 32 kHz by a Neuralynx Digital Lynx system (Neuralynx, USA). LFP signals were obtained by low-pass filtering at 200 Hz and downsampling to 1 kHz. Powerline artifacts were removed by digital notch filtering. The actual spectral data analysis included spectral smoothing that rendered the original notch invisible.

### Data analysis general

All analyses were done in MATLAB (The MathWorks, USA) and using FieldTrip (63) (http://fieldtrip.fcdonders.nl).

### Recording electrodes versus recording sites

During recordings, all ECoG electrodes were referenced against one silver ball implanted epidurally over the other hemisphere. This common reference could lead to artifactual correlations between the signals of separate electrodes. Therefore, all metrics of interaction between distant groups of neurons, that is the pairwise phase consistency (PPC) and Granger causality (GC), were applied after removing the common reference by local bipolar differentiation. That is, the signals from two immediately neighboring electrodes were subtracted from each other. We refer to the ECoG contacts as “electrodes” and to the local bipolar derivations as “recording sites” or just “sites”. All analyses of local neuronal activity used directly the signals recorded from the electrodes, to minimize preprocessing and to minimize reduction in theta amplitude due to theta phase alignment between neighboring electrodes. All analyses of PPC and GC used the bipolar derivations obtained from sites and excluded pairs of sites that shared an electrode.

### Selection of electrodes and sites

The ECoG grids provided dense coverage of dorsal V1, the superficial part of dorsal V2, dorsal V4 and posterior TEO (18, 33). For our analyses, we combined electrodes or sites, respectively, of V1 and V2, and refer to them as V1, and we did the same for V4 and TEO, and refer to them as V4 (see Results for more information on the selection of electrodes and sites, respectively). Monkey K had 45 electrodes on V1, resulting in 40 bipolar sites, and 24 electrodes on V4/TEO, resulting in 19 sites. Monkey P had 72 electrodes on V1, resulting in 64 sites, and 26 electrodes on V4/TEO, resulting in 21 sites.

### Normalization of signals across electrodes and recording sessions

Signal amplitude could vary across electrodes because several separate headstages were used. Furthermore, signal amplitude of a given electrode could vary across sessions, probably due to variable quality of contact to the cortical surface. To equalize the contribution of different electrodes and sessions, we applied a z-transform: Per electrode and session, the raw LFP signal was demeaned and divided by its standard deviation.

### Segmenting data into epochs

Each successfully completed trial contained three periods: The pre-stimulus, the pre-cue and the post-cue period. The pre-stimulus period was the time between fixation onset and stimulus onset. During the pre-stimulus period, monkeys fixated on a fixation point on a gray screen, and there was no stimulus presented and no cue had been nor was presented during that time. The pre-cue period was the time between stimulus onset and cue onset. During the pre-cue period, monkeys kept fixation, the stimuli were continuously present, one tinted yellow the other blue, chosen randomly, and the fixation point had not yet assumed a color, and thereby the attentional cue had not been given. The post-cue period was the time between cue onset and target change. During the post-cue period, monkeys kept fixation, the stimuli were continuously present with their tints and the fixation point was tinted in one of these colors, thereby providing the attentional cue. On approximately half ofF the trials, the post-cue period contained a distracter change, and the data immediately following this event were excluded as explained below.

The pre-stimulus, pre-cue and post-cue periods all were of variable length across trials. The spectral analysis was based on epochs of fixed lengths. Therefore, the described task periods were cut into non-overlapping epochs. We aimed at excluding data soon after events, like stimulus onset, cue onset and distracter change, to minimize effects of post-event transients and non-stationarities on the metrics of rhythmicity and synchronization. Therefore, periods were cut into non-overlapping epochs, starting from the end of the period and stopping, before an epoch would have included data less than 0.5 s after those events. In general, we cut epochs of 1 s length, to achieve a fundamental spectral resolution (Rayleigh frequency) of one Hertz. This was used for the analysis of PPC, GC and phase-amplitude coupling (PAC). The PAC analysis required the prior estimation of the power time course, for which we employed window lengths of ±2.5 cycles per frequency. In this case, epochs were cut such that the power estimation windows excluded data less than 0.5 s after events. The estimation of power spectra was based on 1.6 s epochs, because theta peaks were visible but less conspicuous when 1 s epochs were used. For the time-frequency analysis of power (Fig. S1), epochs of 1 s length were slid over the available post-cue period data in steps of 1 ms.

### Spectral estimation

Epochs were Hann tapered and Fourier transformed. For the PAC analysis, the ±2.5 cycle long windows were also treated in this way. For the analysis of the spatial correlation between theta power and stimulus induced gamma power, the gamma-power estimation used multitaper spectral estimation with seven tapers taken from the discrete prolate spheroidal sequence, defined on 0.5 s long epochs (64).

### Robust regression

We reduced the 1/f^n^ background in power spectra by estimating the 1/f^n^ component and subtracting it. Specifically, for each electrode separately, we pooled all trials of both attention conditions and fitted a line to the log-log power plot between 1 and 10 Hz, using robust regression as implemented in the MATLAB “robustfit” function with default settings. Robust regression uses an iterative procedure that lends less weight to data that are far from the fitted function. Subsequently, the fitted line was subtracted from the power spectrum of each trial, to obtain the power residuals.

### Pairwise phase consistency (PPC) and Phase-amplitude coupling (PAC)

Phase locking was quantified with the pairwise phase consistency (PPC) metric (17). We used PPC both to quantify the locking between LFPs recorded from separate sites, the locking between microsaccades and LFP, and the locking between the LFP phase and its amplitude fluctuations, that is, the PAC (phase-amplitude coupling) (65). PPC is not biased by the number of epochs, whereas the more conventional coherence metric has that bias. Essentially, the PPC calculation proceeds in two steps. First, the relative phases are calculated for the multiple epochs of the two signals. The second step is the crucial step: In conventional coherence calculation, those relative phases are averaged, which leads to the bias by epoch number; in PPC calculation, all possible pairs of relative phases are formed, the cosines between those relative phases are determined and those cosine values are averaged.

To quantify PAC, we computed the PPC between the LFP at lower frequencies, the “phase-frequencies”, and the time-varying power at higher frequencies, the “amplitude-frequencies”. One-second long epochs of the raw LFP and of its time-varying power were Fourier transformed, and locking among the phase estimates at the phase-frequencies was quantified as the PPC across all available epochs. PAC can in general only be estimated for pairs of phase- and amplitude-frequencies, for which the amplitude frequency is higher than the phase frequency. In addition, the estimation of time-varying power entails low-pass filtering, because windows of some length in time are required to estimate the power with some specificity in frequency. Consequently, PAC can only be estimated for pairs of phase- and amplitude-frequencies, for which this low-pass frequency is above the phase frequency. Power is estimated on the basis of epochs and tapers of finite length. As described above, we chose epochs of ±2.5 cycle length per frequency. In order to assess the resulting low-pass filtering, we applied the power estimation 10000 times to a random Gaussian process of the same length as the data epochs, and determined the frequency, at which this low-pass filtering reduced the average power to less than 70% of the power in the passband. For example, for 50 Hz, this cutoff frequency was 7.7 Hz. This procedure was applied for each amplitude frequency, and the PAC for this amplitude frequency was only considered up to the respective phase frequency. The excluded combinations of phase-frequencies and amplitude-frequencies are masked with black in the figures. The PAC results shown here use phase and power estimates from the same electrode. We also calculated PAC by combining phase estimates from one electrode with power estimates of neighboring electrodes, and this left the results essentially unchanged.

### Granger causality

We used the non-parametric estimation of Granger causality (GC) (66). For this, Fourier spectra were estimated as described above and entered into a non-parametric spectral matrix factorization (NPSF) as implemented in the FieldTrip toolbox (63).

### Statistical testing

The confidence intervals shown for power and PPC spectra in Figure 1 were estimated with a bootstrap procedure (1000 bootstrap replications for power, 500 for PPC) (67): Spectra were first averaged across electrodes (for power) or site pairs (for PPC), and subsequently, the bootstrap was performed across epochs. All statistical comparisons were based on non-parametric permutation and included corrections for the multiple comparisons made across frequencies. We illustrate the procedure for the comparison of power between the two attention conditions. The power difference between the attention conditions was first averaged over all electrodes per monkey and then over the two animals, giving the observed power difference per frequency. Subsequently, the following procedure was done 1000 times: 1) The attention conditions were randomly exchanged between epochs, keeping the original number of epochs per attention conditions constant; 2) The average power difference was calculated as described for the observed data; 3) The maximal (minimal) difference across all frequencies was placed into the randomization distribution of maximal (minimal) values; 4) The 2.5^th^ percentile of the minimal values and the 97.5^th^ percentile of the maximal values were taken as statistical thresholds. The observed differences were compared to those thresholds. This procedure implements a non-parametric version of a two-sided test with multiple comparison correction (68). The same procedure was used for comparing power, PPC, GC and PAC values between attention conditions; for power and PAC, we used 1000 permutations, for PPC and GC 500 permutations.

The spatial correlation coefficients and the PAC values were tested in two ways: They were compared between attention conditions as described, and they were additionally tested for the presence of significant correlation or PAC. In the case of PAC, the comparison was done between the observed values and a randomization distribution obtained by randomly pairing raw LFP epochs and power time courses 1000 times. After each random pairing and recalculation of PAC, maximal and minimal values across all frequency-frequency pairs were placed into the respective randomization distribution, and further testing proceeded as described. In the case of the spatial correlations, the comparison was done between the observed values and zero, because the Spearman rank correlation has no bias; the randomization was done by randomly pairing electrodes between the theta power residuals and the stimulus induced gamma. After each randomization, maximal and minimal correlation values across all tested frequencies were placed into the respective randomization distribution, and further testing proceeded as described.

### Microsaccade detection

Raw vertical and horizontal eye position signals were low-pass filtered by replacing each value with the average over itself ±15 samples (at 1 kHz sampling rate). Signals were then differentiated in time to obtain the vertical and horizontal velocities. Those were combined to obtain the eye speed irrespective of the direction of eye movement. Per trial, the standard deviation of eye speed was determined, and any deviation larger than 5 SDs and lasting for at least 30 ms was considered a saccadic eye movement. Saccadic eye movements that remained within the fixation window were considered microsaccades (MSs).

### Stratification

We intended to test whether some of the observed differences were due to differences in the rate of MSs or in the power of theta, which existed between attention conditions. To this end, we used a stratification approach, that is, we randomly subsampled the available data to equate as well as possible the distributions of MS rates or theta power (69). For MS stratification, we first calculated MS density by convolving the MS sequence with a Gaussian kernel with an SD of 150 ms (truncated at ±500 ms). For each epoch, we calculated the average MS density, which was then used for stratification. For theta power stratification, we estimated and removed the 1/f^n^ component for each electrode, averaged over electrodes, and used the resulting average residual theta (3-5 Hz) power for stratification. We describe the stratification procedure for a given parameter (MS density or theta power): The parameter distributions were compiled for the two attention conditions and binned into 40 equally spaced bins. For each bin, the number of entries for the two attention conditions was equated by random subsampling with a procedure that aims at equating the parameter averages between the conditions as well as possible. This procedure is applied to the distributions per bin: 1) The condition with more entries is defined as the larger condition, the other as the smaller condition; 2) The mean of the parameter for the smaller condition is calculated and taken as target value; 3) The larger condition is randomly subsampled, by first drawing one entry at random, and then proceeding as follows: a) A further entry is randomly drawn; b) If the mean of the current bin entries (or the starter entry) is smaller (larger) than the target value, the new entry is added if it is larger (smaller), otherwise it is discarded and a new random draw is performed. This latter step aims at equating means; if no such entry is present, a randomly drawn entry is accepted.

## Acknowledgements

PF acknowledges grant support by DFG (SPP 1665, FOR 1847, FR2557/5-1-CORNET), EU (HEALTH-F2-2008-200728-BrainSynch, FP7-604102-HBP, FP7-600730-Magnetrodes), a European Young Investigator Award, NIH (1U54MH091657-WU-Minn-Consortium-HCP), and LOEWE (NeFF). The authors thank Jarrod Dowdall for help with microsaccade detection and Martin Vinck for helpful comments on the manuscript.

## Supporting Information

### Supplementary Figure Legends

**Fig. S1.**
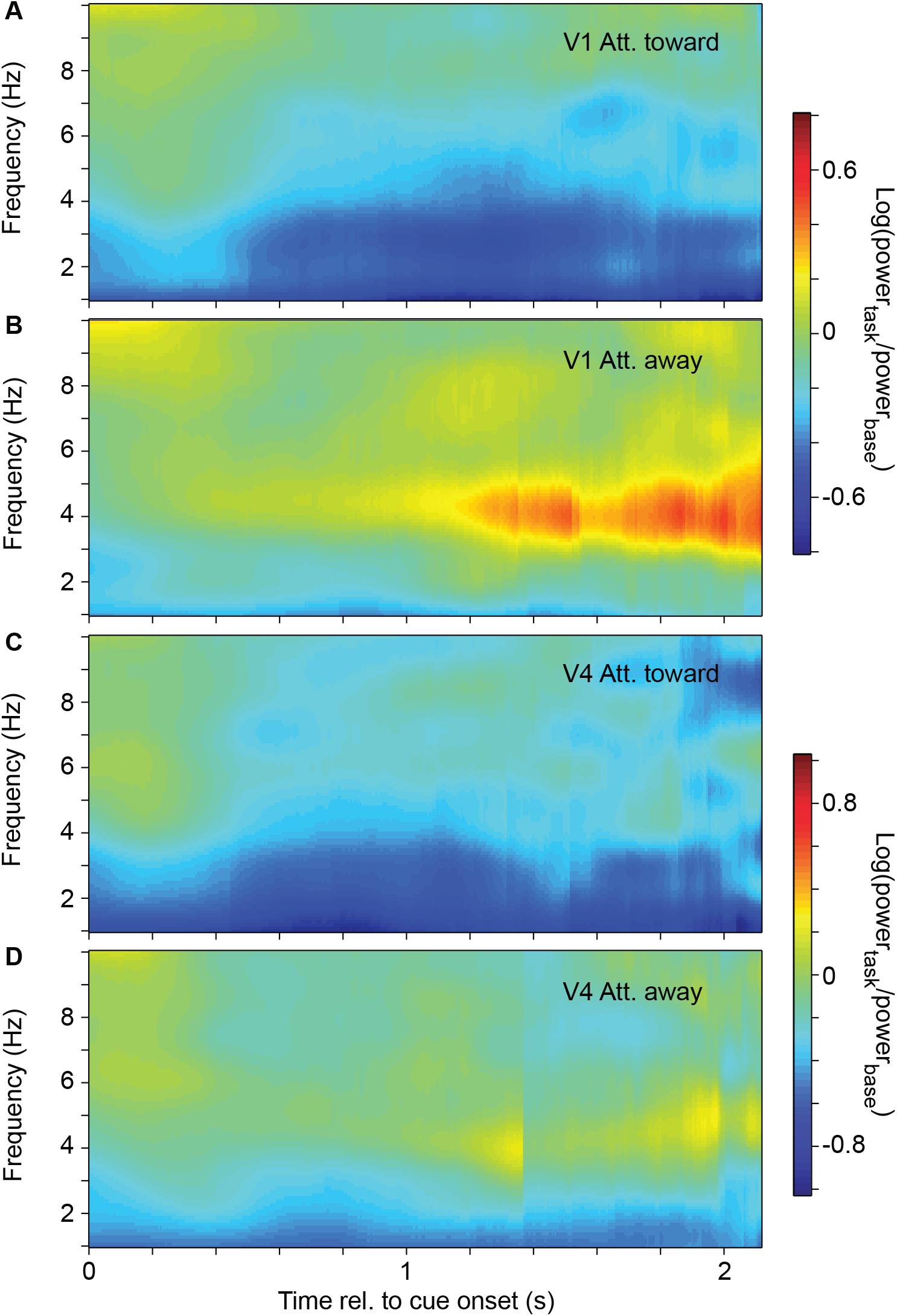
Average time-frequency analysis of power. (*A,B*) Time-frequency power averaged over all selected V1 electrodes of both monkeys, with attention toward (*A*) and away from (*B*) the activating stimulus. Power is expressed as the logarithm (base 10) of the ratio between the power during stimulation plus task performance, and the power during the pre-stimulus baseline. (*C,D*) Same as (*A,B*), but for area V4. Note that these time-frequency power analyses are based on trials of variable length. As a consequence, the power estimates for later time points are based on fewer trials. The dropping out of trials causes the visible temporal discontinuities (particularly notable for V4, attend away). The log-ratio of power is not biased by the number of trials, and therefore, the visible increase of theta power in V1 with attention away is most likely genuine.

**Fig. S2.**
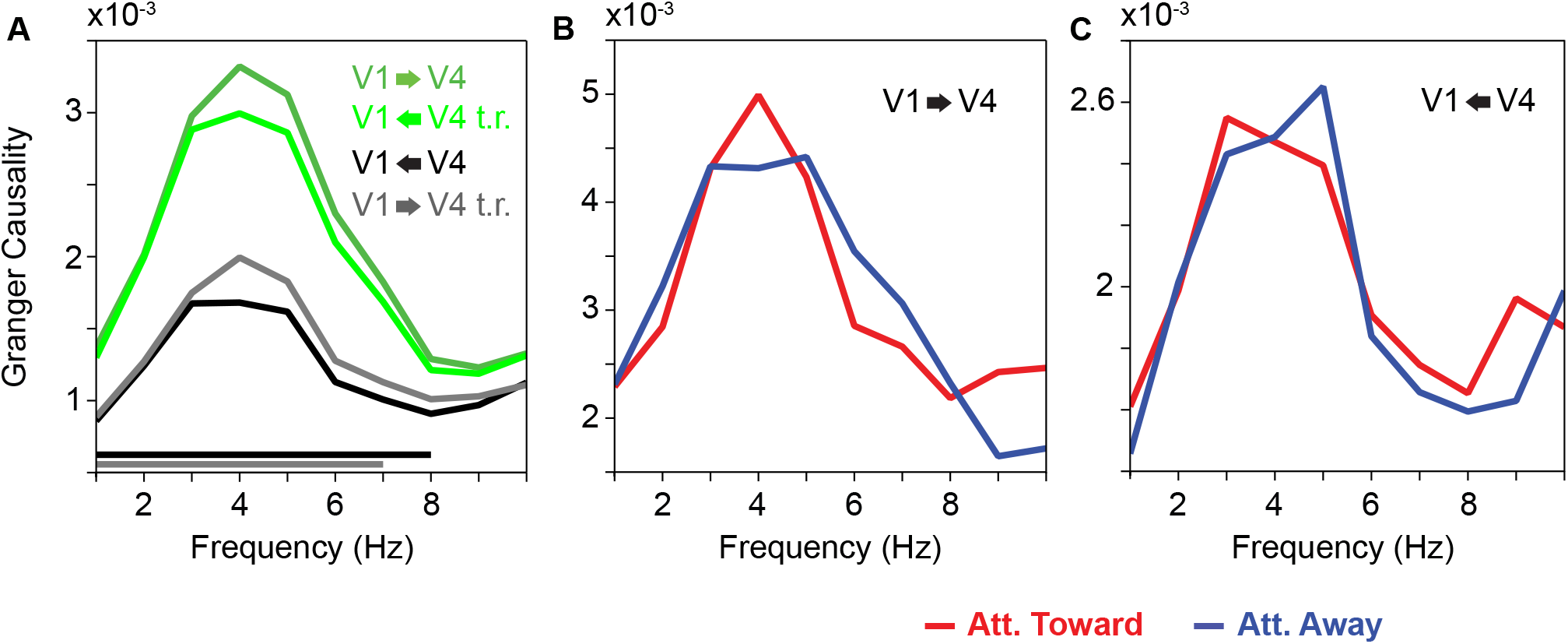
Average low-frequency Granger causality (GC) spectra between V1 and V4 sites, controlling for differences in noise between areas or for theta power differences between attention conditions. (*A*) Average GC-influence spectra between V1 and V4 in the feedforward and feedback directions for the original (dark green and black respectively) and time-reversed signals (gray and light green respectively). Lines on the bottom indicate frequencies with a significant difference between bottom-up and top-down for the original (black) and the time-reversed signals (gray) (p<0.05; non-parametric permutation test with correction for multiple comparisons across frequencies). (B) Average GC-influence spectra between V1 and V4 in the feedforward direction after theta power stratification, with attention toward (red) and away from (blue) the activating stimulus. (*C*) Same as *B*, but for the feedback direction. Neither *B* nor *C* revealed any significant differences.

**Fig. S3.**
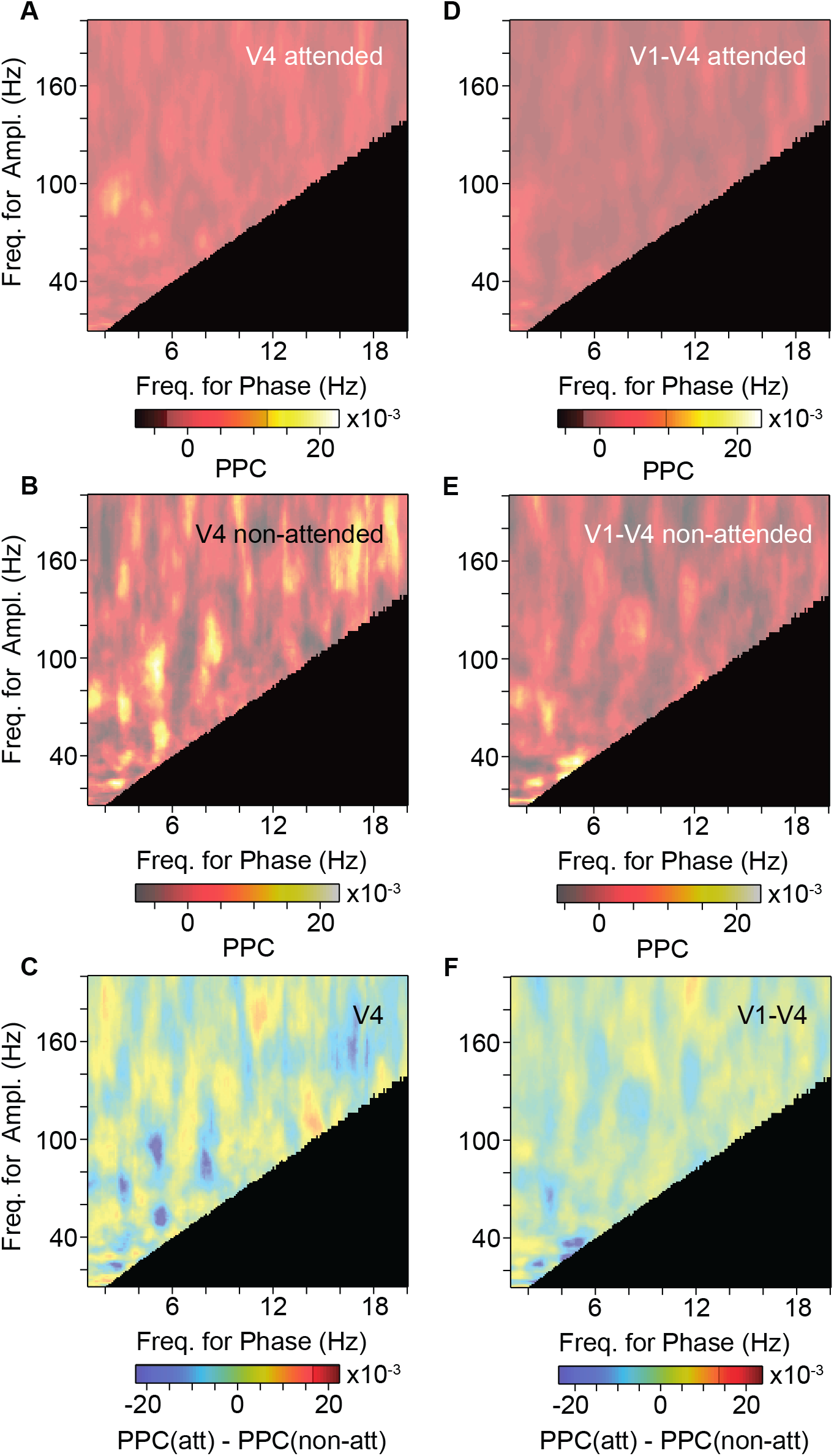
Modulation of PAC by selective attention, excluding epochs with microsaccades. (*A, B*) Average PAC in area V4 with attention toward (*A*) and away from (*B*) the activating stimulus. (*C*) Average PAC difference in area V4 between the two attention conditions shown in *A* and *B*. (*D*), (*E*), (*F*) Same as *A, B, C*, but using the phase in area V1 and the amplitude in area V4. The semitransparent gray mask indicates frequency pairs with non-significant PAC, or PAC difference, respectively (p<0.05; non-parametric permutation test with correction for multiple comparisons across frequency pairs). The black area indicates frequency pairs excluded from the analysis (see Materials and Methods).

**Fig. S4.**
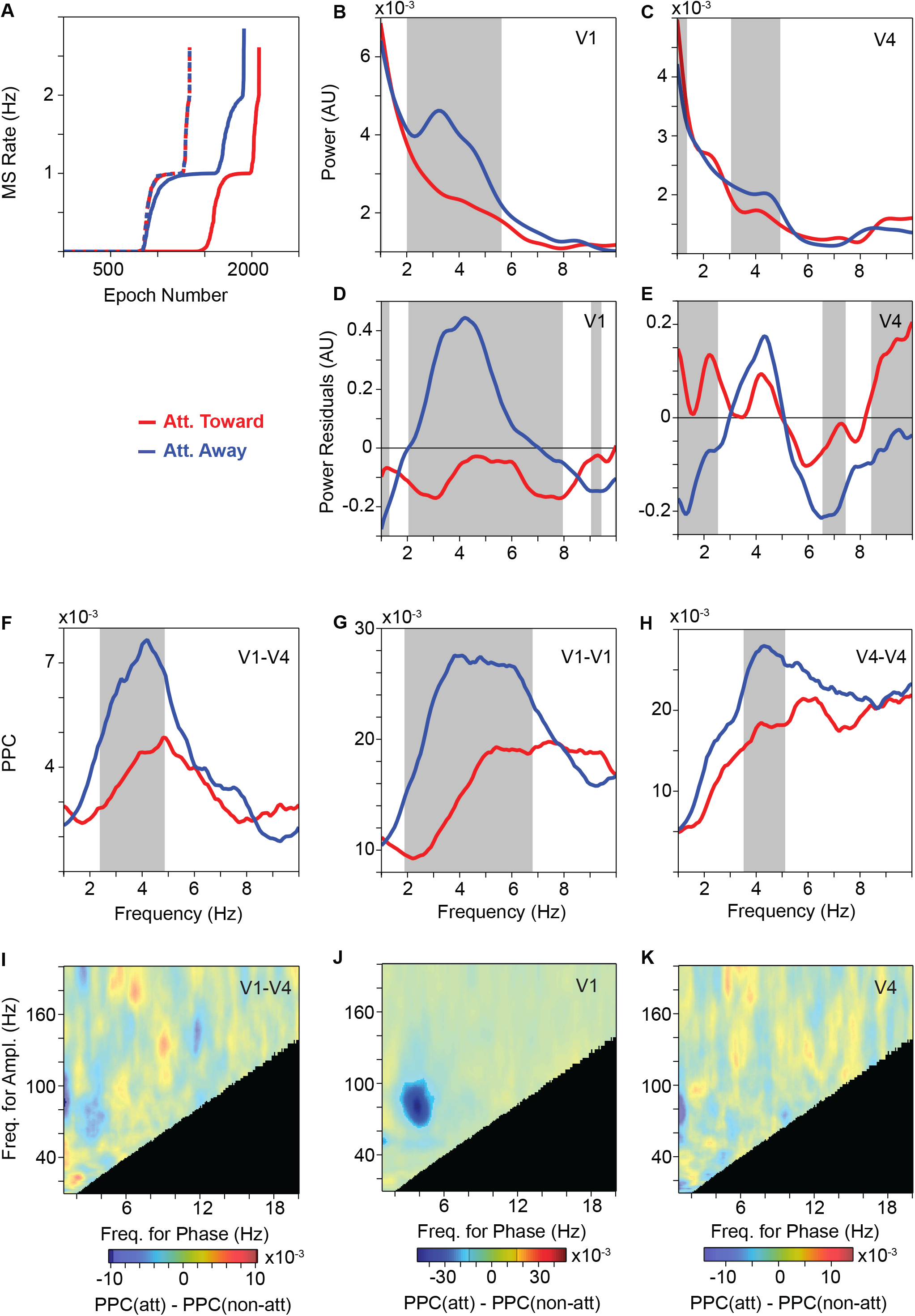
Attention contrast, controlled for microsaccade (MS) rate. (*A*) Cumulative distribution of MS rate with attention toward (red) and away (blue) from the activating stimulus. Solid lines show data before stratification; dashed lines show data after stratification. Note that after stratification, the lines for the two attention conditions overlap essentially perfectly. (*B*) Average LFP power spectra in area V1, with attention toward (red) and away (blue) from the activating stimulus. (*C*) Same as *B*, but for area V4. (*D*), (*E*) Same as *B* and *C*, after removing the 1/f^n^ component. (*F*) Average LFP phase locking between sites in area V1 and sites in area V4, with attention toward (red) and away from (blue) the activating stimulus. (*G*), (*H*) Same as *F*, but for pairs of sites within area V1 (*G*) and area V4 (*H*). (*B-H*) Gray-shaded regions indicate frequencies with a significant difference between attention conditions (p<0.05; non-parametric permutation test with correction for multiple comparisons across frequencies). (*I*) Average PAC difference between the two attention conditions using the phase in area V1 and the amplitude in area V4. (*J*) Same as *I*, but for area V1. (*K*) Same as *I*, but for area V4. The semitransparent gray mask indicates frequency pairs with non-significant PAC difference (p<0.05; non-parametric permutation test with correction for multiple comparisons across frequency pairs). The black area indicates frequency pairs excluded from the analysis (see Materials and Methods).

**Fig. S5.**
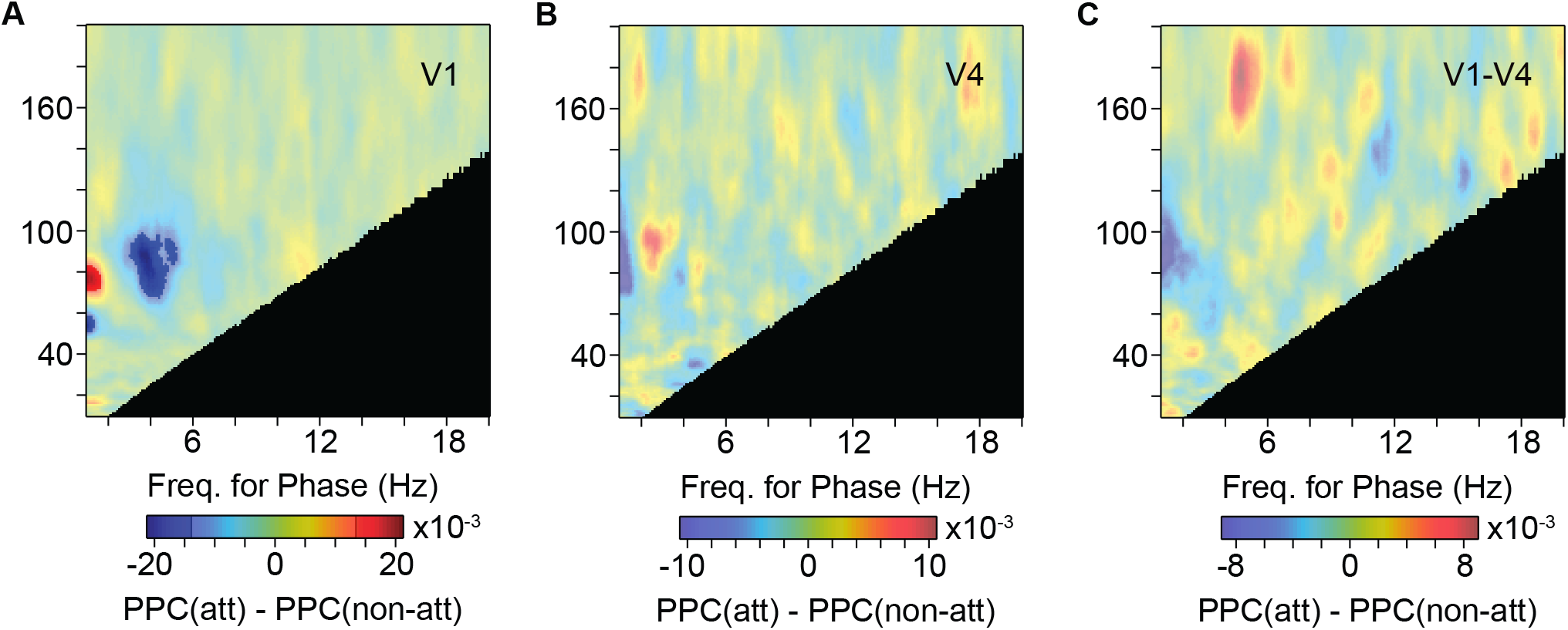
Modulation of PAC by selective attention, after theta power stratification. (*A*) Average PAC difference between the two attention conditions in area V1. (*B*) Same as *A*, but for area V4. (*C*) Same as *A*, but using the phase in area V1 and the amplitude in area V4. The semitransparent gray mask indicates frequency pairs with non-significant PAC difference (p<0.05; non-parametric permutation test with correction for multiple comparisons across frequency pairs). The black area indicates frequency pairs excluded from the analysis (see Materials and Methods).

**Fig. S6.**
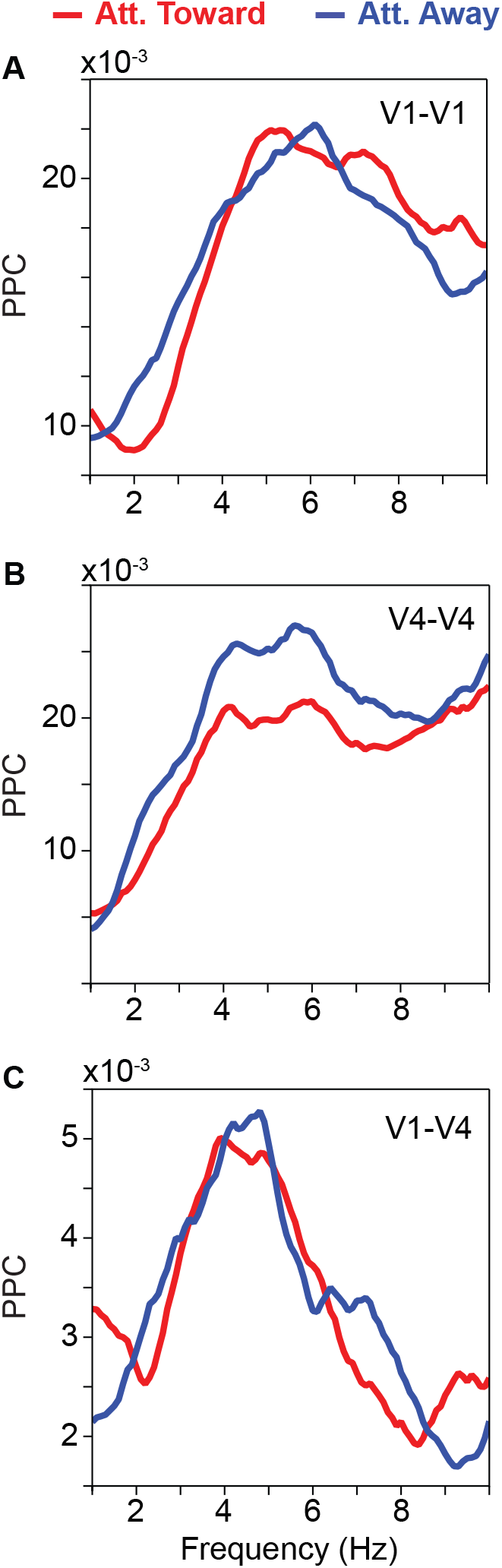
Average low-frequency LFP phase-locking (PPC) spectra and their modulation by selective attention, after theta power stratification. (*A*) Average LFP phase locking between sites within area V1 with attention toward (red) and away from (blue) the activating stimulus. (*B*) Same as *A*, but between sites within area V4. (*C*) Same as *A*, but between sites in area V1 and sites in area V4. Neither *A, B* nor *C* revealed any significant differences.

